# From Cell-Lines to Cancer Patients: Personalized Drug Synergy Prediction

**DOI:** 10.1101/2023.02.13.528276

**Authors:** Halil Ibrahim Kuru, A. Ercument Cicek, Oznur Tastan

## Abstract

Combination drug therapies are effective treatments for cancer. However, the genetic heterogeneity of the patients and exponentially large space of drug pairings pose significant challenges for finding the right combination for a specific patient. Current *in silico* prediction methods can be instrumental in reducing the vast number of candidate drug combinations. However, existing powerful methods are trained with cancer cell line gene expression data, which limits their applicability in clinical settings. While synergy measurements on cell lines models are available at large scale, patient-derived samples are too few to train a complex model. On the other hand, patient-specific single-drug response data are relatively more available. In this work, we propose a deep learning framework, <u>P</u>ersonalized <u>D</u>eep <u>S</u>ynergy <u>P</u>redictor (PDSP), that enables us to use the patient-specific single drug response data for customizing patient drug synergy predictions. PDSP is first trained to learn synergy scores of drug pairs and their single drug presonses for a given cell line using drug structures and large scale cell line gene expression data. Then, the model is fine-tuned for patients with their patient gene expression data and associated single drug response measured on the patient *ex vivo* samples. In this study, we evaluate PDSP on data from three leukemia patients and observe that it improves the prediction accuracy by 27% compared to models trained on cancer cell line data. PDSP is built and available at https://github.com/hikuru/PDSP

## 1 Introduction

Combination therapies, in which multiple drugs are administered concurrently, have emerged as alternatives to single drug therapies for various complex diseases including cancer [1, 18, 20, 23, 26], diabetes [15], human immunodeficiency virus [8], and SARS-CoV-2 [32]. These therapies are reported to provide more effective treatment [9, 29], overcome drug resistance [13], and decrease side effects by enabling reduced doses [1, 38]. While combination therapies confer benefits to cancer patients compared to single drug ones, personalizing drug combinations for individual patients is challenging. The effectiveness of drug combinations varies across patients due to differences in their genotypic and phenotypic variation [10, 27].

To facilitate drug combination discovery, preclinical studies rely on screening drug combinations in cancer cell lines experimentally. However, even for medium-sized drug panels, exhaustive experimentation with all possible combinations is impossible as the space of possible combinations increases exponentially with the number of drugs [11, 24, 30]. To aid the combination discovery efforts, though most are not personalized, many *in slico* methods have been proposed [4, 25]. Various mathematical modeling techniques [39], network-based models [7], and classical machine learning techniques have been applied to predict drug synergy [2, 17, 21, 33]. These models have been trained on various types of data, including drug chemical properties, cell line untreated and/or treated gene expression [34], drug-target interaction [2], and protein-protein interaction networks [7]. Yet, the prediction performance was limited due to limited training data.

Over the last decade, the research community focused on compiling large scale drug synergy datasets and organized several challenes to develop more advanced methods to exploit these resources [2, 35]. For instance, AstraZeneca and Sanger [35] released more than ten thousand experimentally tested drug combinations, where cell viability was measured across various cell lines. Specifically, this DREAM challenge considered 11, 759 drug combination screenings that span 85 compounds and 85 cell lines. Similarly, NCI-ALMANAC study[14], evaluated the combination effects of 104 FDA-approved cancer drugs on 60 cell lines. This dataset contains nearly 300,000 drug pair-cell line combinations.

With the availability of large-scale training data, we [19] and others [17, 22, 28, 31, 37] have developed deep learning based solutions to identify synergistic drug combinations. Preuer et al. presented DeepSynergy, which takes the chemical structures and cancer cell line gene expression data as input to a fully-connected neural network to predict synergy of drug pairings. MatchMaker [19] uses a multi-modal architecture and employs three fully-connected neural network modules to learn cell line-specific representations of drugs and predicts drug combination synergy scores. Finally, in a recent approach, DeepDDS [31] uses a graph convolutional neural network to predict combination synergy scores. This approach represents molecules as graphs, where atoms correspond to nodes and bonds correspond to edges.

While the deep learning models showed superior performance in predicting synergistic drug pairs, two critical challenges stand in the way of deploying deep learning models to personalize predictions for cancer patients. Cancer cell line expression data, available for many cancer cell lines, has been used to contextualize the synergy effects in these prior works. However, cell line expressions are only a proxy for the patient expression data; and do not capture the individual patient variation. The models require a large amount of training data to learn to patient synergy scores, but measurements on patient-derived samples are not available at large. To tackle patient customized synergy prediction problem, He et al. proposed a computational-experimental setup where they use *in silico* prioritization and *ex vivo* testing in patient-derived samples to identify customized synergistic combinations for individual cancer patients. Since the available patient response data is small, they build a random forest model to learn single-drug dose-response curves of patients and use this model to predict drug combinations. Unlike the above-mentioned deep learning models, the model does not benefit from the available large-scale synergy measurements obtained on cell lines. To the best of our knowledge, no model can make use of both the available large-scale cell line data and the limited patient data to achieve patient-level drug synergy prediction. In this work, we address this problem.

To tackle the patient-specific synergy prediction task, we propose a deep learning-based solution (Figure 1). In the first stage, we train the model using the large-scale single drug response and synergy scores of drug pairs measured on cell lines (Figure 1.1). The drug response prediction module enables us later to calibrate our model with the limited single drug measurements on patient-derived samples. In the second stage, we fine-tune the models using gene expression and drug response data measured on patient-derived *ex vivo* samples. To customize the models for each patient, we present two novel patient fine-tuning strategies that make use of the patient single drug response data. (i) we use the pre-trained model and then fine-tune the model with drug sensitivity labels of all training patients data (Figure 1.2.a); (ii) We use patient-specific fine-tuning, in which we fine-tune the pre-trained model for each patient separately (Figure 1.2.b). We then predict test patients’ synergy data with the corresponding models (Figures 1.3.a and 1.3.b). We demonstrate that these strategies accurately predict synergy for the patients while maintaining the model’s comparable performance on cell lines. Currently, there is no comparable study that uses deep learning models with measurements on patient data; therefore, this model represents an advance toward personalizing drug synergy predictions.

**Fig. 1.**
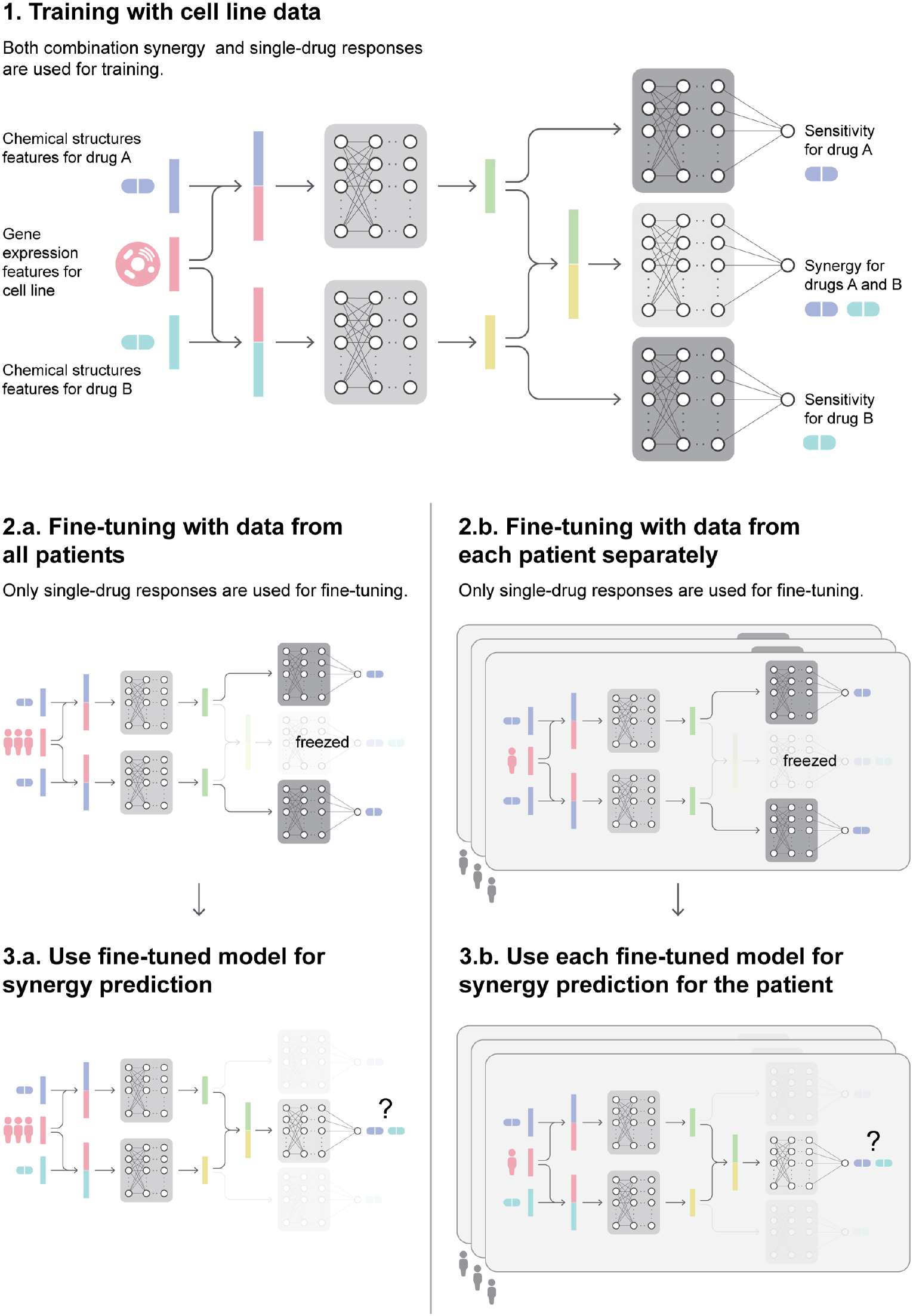
The general pipeline for PDSP framework. 1) The architecture and the training procedure of the PDSP model. The chemical features of each drug are concatenated with gene expression features of the cancer cell line and input to drug encoder networks. They learn a representation of each drug separately conditioned on cancer cell line gene expressions. The synergy aggregations network takes both drugs’ representations and applies fully connected layers to predict synergy scores. The drug decoder network takes conditioned drug representations and applies connected layers to predict the drug sensitivities. 2) Fine-tuning the trained PDSP model. The pre-trained PDSP model is used for fine-tuning the drug encoders by using patient single-drug responses and gene expressions for all patients (2.a) and per patient (2.b). 3) Synergy testing with respective fine-tuned models from the previous part.

## 2 Methodology

For a drug pair < *i, j* > and a cell line *k*, we aim to predict pair synergy scores *y_i,j,k_* ∈ ℝ along with each drug’s sensitivities in cell line *k*. We binarize the individual compound sensitivity scores (see Section 2.1). Let *s_i,k_* ∈ {0,1}, and *s_j,k_* ∈ {0,1} represent drug *i* and *j*’s individual sensitivity classes, where 0 indicates resistant and 1 indicates sensitive. We represent each drug with its chemical features *d_i_* and *d_j_*, and cancer cell lines with gene expression profiles *c_k_*. The model is trained jointly to predict the two drugs’ individual sensitivity classes and pair synergy score by using drug and cell line feature vectors (*d_i_*, *d_j_* and *c_k_*).

### 2.1 Datasets

#### Cancer cell line drug response dataset

We use the Bliss synergy scores measured on cancer cell lines for synergy measurements and half-maximal inhibitory concentration (*IC*_50_) values for sensitivity measurements. We obtain the data from the DrugComb [36] database through https://drugcomb.fimm.fi/ (version v1.5, downloaded on Feb 2022). We binarize *IC*_50_ values as sensitive and resistant. Following Chang et al., we designate cell lines as sensitive to a drug if the natural logarithm of its *IC*_50_ is less than −2 and resistant otherwise. Since DrugComb includes data from multiple sources, it sometimes contains replicates for the same drug pair. In these cases, we average Bliss scores across replicate measurements. Not all cancer cell line gene expression profiles were available at Iorio et al., so we filtered out cell lines for which gene expression profiles were not available. Our final dataset contains 330,103 samples with 3,068 drugs and 81 cancer cell lines.

#### Patient drug response and synergy dataset

We obtain patient synergy and sensitivity data from [12]. This work provides 654 sensitivity measurements on patient-derived samples covering 218 compounds on*ex vivo* and three patients. He et al. has also made drug combination experiments and reported 20 experimentally verified drug combinations results, of which 10 are synergistic, and the remaining 10 are antagonistic. These 20 combinations include 16 drugs for the three patients. We use the patient drug response data for finetuning the models and use the synergy measurements for evaluating the final model for personalized synergy prediction.

#### Model features

Our model relies on two sets of features, one set for representing drugs and the other set for cell line representation. We represent drugs with their chemical structure features, which we calculate using ChemoPy Python library [5] (more information provided in our earlier work [19]). Cell lines are represented with their untreated gene expression profiles [16], and RMA normalized profiles are obtained from https://www.cancerrxgene.org (downloaded on Dec 2019). Since we apply our model to patient expression profiles from He et al., we use available gene expression profiles from this study. To be able to apply PDSP on patient gene expression profiles, we take the intersection of 17, 737 genes in cancer cell lines and 634 genes in patient data. The intersection of these two gene sets contains 542 genes; we use expressions of these genes as gene expression features. The cancer cell line gene expressions and the patient gene expressions are measured under different conditions. We apply quantile normalization [3]. We input two drugs *d_i_,d_j_* ∈ ℝ^541^ and a cell line *c_k_* ∈ ℝ^542^ to our model and predict *y_i,j,k_* ∈ ℝ, *s_i,k_* ∈ ℝ, and *s_j,k_* ∈ ℝ simultaneously.

### 2.2 Training Loss

We train the model with a loss function that combines the regression loss *L_reg_* and the classification loss *L_cls_*:

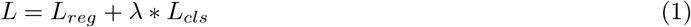

where λ is a hyperparameter that adjusts the weight between regression and classification losses. We adopt *mean-squared-error* as the regression loss, and *weighted binary cross entropy* (wBCE) as the classification loss:

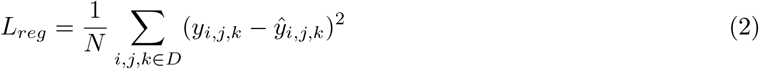

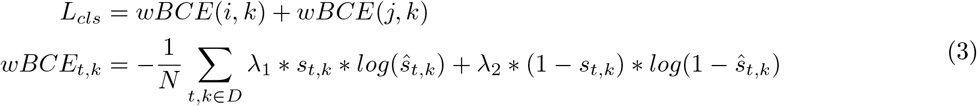

where *N* is the number of drug combinations in training dataset *D. y_i,j,k_* is the ground-truth Bliss score of drug combination <*i,j*> on cell line *k. s_t,k_* is the ground-truth sensitivity class of drug *t* on cell line *k. ŷ_i,j,k_* and *ŝ_t,k_* are the corresponding predictions. λ_1_ and λ_2_ are the loss weights for sensitive and resistant drugs on cell line *k*.

### 2.3 The Model Architecture

PDSP model builds upon our previous work Matchmaker [19]. Figure 1 provides an overview of the PDSP architecture. We first represent cell lines and two drugs as a collection of vectors *d_i_, d_j_*, and *c_k_*. With these vector representations, PDSP aggregates both cancer cell line and drugs’ information and predicts drug sensitivities and combination synergy in two stages. In the first stage, PDSP encodes drug and cancer cell line information with Drug Encoder Networks (DEN). DENs take both drug and cancer cell line representations and learn conditional latent representations of the drugs. We force both DENs to share the same architecture and model weights to get order-invariant latent representations. In the second stage, PDSP uses these conditional encodings to predict sensitivity classes and combination synergy score. This second stage contains two Drug Decoder Networks (DDN) and one Synergy Aggregation Network (SAN). DDN takes encoded conditional latent representation and decodes it to the drug’s sensitivity on a given cell line. Similarly to DENs, DDNs share the same architectures and model weights. While drug encoders learn representations of each drug, they do not provide information about their combinations in a cell line. Therefore, SAN first aggregates drugs’ latent representation by concatenation and provides the combination information. Finally, an aggregated representation is used to predict drug combination synergy.

#### Drug Encoder Network (DEN)

The PDSP model consists of two identical drug encoders, *DEN_i_* and *DEN_j_*. These encode the cancer cell line features (*c_k_*) and two drugs’ features (*d_i_* and *d_j_*) into a conditional latent space. The DEN model takes gene expression data and concatenates it with the features of each drug in the combination. This concatenated information is then passed through a series of fully-connected (FC) layers. Each FC layer is followed by a non-linear activation function (ReLU), except for the last FC layer and dropout layers. The formulation of DEN is as follows:

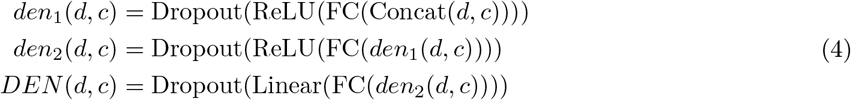

where *d* and *c* denotes the drug and cell line features, *den*_1_ and *den*_2_ are the intermediate layers. The *DEN_i_* and *DEN_j_* encoders are designed to have the same model parameters and weights, which allows for the same latent representations to be achieved for the two drugs (order-invariance). This shared structure enables the model to learn the relationships between the drugs and cancer cell lines consistently, regardless of the order in which the drugs are presented. The outputs of the encoders, *h_i,k_* and *h_j,k_* ∈ ℝ^*m*^, are multi-dimensional representations of the drugs in the conditional latent space. We use these drug encodings as inputs to downstream tasks, drug sensitivity, and drug combination prediction. *DEN_i_* and *DEN_j_* provide drugs encodings *h_i,k_* and *h_j,k_* ∈ ℝ^*m*^ as follows:

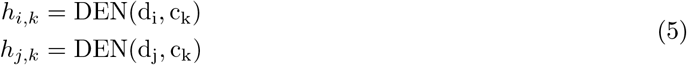

#### Drug Decoder Network (DDN)

Based on the latent drug encodings *h_i,k_* and *h_j,k_*, we train two identical drug decoder networks *DDN_i_*, and *DDN_j_* to output drug sensitivities *ŝ_i,k_* and *ŝ_j,k_*. DDN uses a feed-forward neural network architecture with three FC layers. The first two FC layers apply and ReLU activation and dropout, respectively. Finally, a final FC layer with a linear activation function is used to obtain the drug sensitivity status. The DNN formulation is given below:

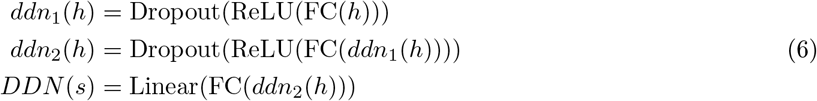

where *h* corresponds to the conditional drug encodings (*h_i,k_* or *h_j,k_*), *ddn*_1_ and *ddn*_2_ are the intermediate layers. Both DDNs use the same model weights, and sensitivity predictions *ŝ_i,k_* and *ŝ_j,k_* for drugs *i* and *j* on cancer cell line *k* is calculated as follows:

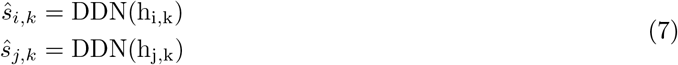

#### Synergy Aggregation Network (SAN)

Similarly to DDN, synergy aggregation network uses feedforward neural network architecture. SAN takes latent drug representations *h_i,k_* and *h_j,k_* and predicts drug combination synergy *ŷ_i,j,k_*. Since the inputs of SAN is a drug combination, it first concatenates drug features, then applies similar network architecture to DDN. We apply linear activation at the last layer of SAN since it outputs the Bliss synergy score. The SAN model is given below:

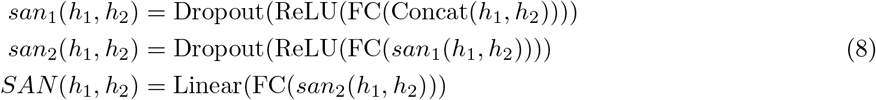

where *h*_1_ and *h*_2_ denotes conditional drug encodings in the combination, *san*_1_ and *san*_2_ are the intermediate layers. With these formulations, we obtain predicted synergy score as *ŷ_i,j,k_* = *SAN*(*h_i,k_, h_j,k_*).

### 2.4 Customizing for Patients

We initially train PDSP with drug combination data derived from cancer cell line experiments. In this pretrained model, we freeze SAN and fine-tune the model with patient single-drug responses. Fine-tuning provides us to calibrate DENs to the patient-derived sample data. We apply two patient fine-tuning strategies and asses their predictive performance.

We denote the initial trained model weights on cancer cell line as *Φ_c_*, patients’ gene expression profiles *c_pt_* ∈ ℝ^542^ (*t* ∈ {1,2,3}), drugs’ chemical features as *d_i_* ∈ ℝ^541^ and fine-tune *Φ_c_* as follows:

#### Strategy 1: Fine-tune model with all patients

We use *Φ_c_* as the pre-trained model weights for PDSP. We feed patients’ gene expression profiles and drugs’ chemical structure features into the model and run the PDSP training pipeline for single-drug responses. After fine-tuning, we use this new model weights *Φ_c,all_* for synergy prediction of all patients. We refer to this model as PDSP^st1^. This strategy is useful when we have a set of single-drug responses for multiple patients.

#### Strategy 2: Fine-tune model for each patient

In the second strategy, we fine-tune pre-trained weights *Φ_c_* for each patient separately. This lead to *n* different models for each of the *n* patients: *Φ_c,t_* (*t* ∈ {1, …*n*}. We evaluate each patient’s drug combination synergy score with the corresponding model’s weights *Φ_c,t_*. We refer to the models generated using this strategy PDSP^st2^.

## 3 Results

### 3.1 Experimental Setup

We follow Preuer et al. for performance benchmark and use *leave-drug-combination-out* approach. We divide the whole drug combination data into train, validation, and test splits by considering that a drug pair <*i, j* > can only be involved in one of the splits. The ratios of train, validation and test splits are 60%, 20%, and 20%, respectively. We tune the model hyperparameters based on the performance of the validation set. We use these models to report performance on cancer cell line test data. We also use the best PDSP model weights *Φ_c_* as the pre-trained model for patient data calibration as explained in 2.4.

### 3.2 Performance Benchmark

#### Regression Performance on Cancer Cell Lines

We compare PDSP’s performance to the state-of-the-art models: MatchMaker [19], DeepSynergy [28], TreeCombo [17] and DeepDDS [31]. We also use a standard machine learning algorithm, Random Forest, as the baseline model.We evaluate them based on MSE, Pearson, and Spearman rank correlation coefficients between the actual Bliss synergy scores and the predicted scores on the unseen cell line test set. Table 1 shows the model performances for these performance metrics. Among these methods, PDSP and MatchMaker achieve the top performances. Both models show comparable performances across all metrics. While MatchMaker shows slightly better performance on MSE (41.60 vs. 41.95), PDSP is better on the Spearman correlation coefficient (64% vs. 65%). They achieve the same performance in the Pearson correlation coefficient: 91%. These results show that adding drug response prediction nodes to enable patient calibration did not sacrifice on the synergy prediction performance; PDSP achieves on par synergy prediction performance on cell-lines with MatchMaker (i.e., no patient calibration). When we compare PDSP’s performance with the remaining models, PDSP achieves ~ 20% MSE and 2% to 7% correlation improvements over the next best model, DeepSynergy. DeepDDS, a model that uses graph neural networks to encode drug chemical structure, performs worse than the baseline method, Random Forest.

**Table 1.**
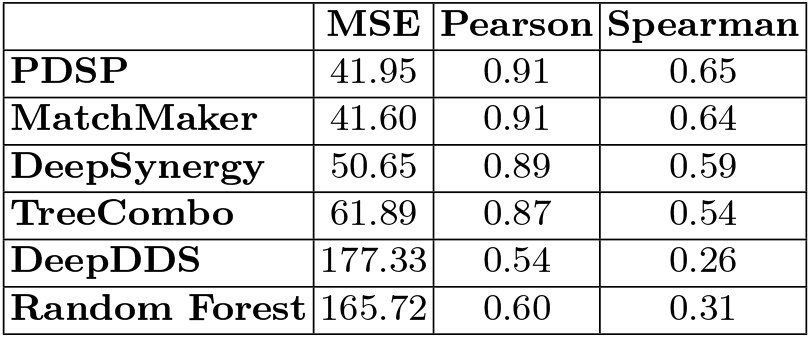
Comparison of synery predictions of the models based on MSE, Pearson and Spearman correlation coefficients.

We further assess the cell line specific performances of the models by measuring the regression performance metrics for each cell line separately. We provide the list of 10 cell lines for which our method performed the best. We also provide the number of instances that cell line is observed in training data in Supplementary Tables 1-3. We observe that the number of cases does not have a substantial effect on the performance.

#### Classification Performance on Cancer Cell Lines

We also the classification performance of the synergy model. We binarize the actual Bliss scores: synergistic pairs have positive Bliss scores, and antagonistic pairs have negative Bliss scores. We perform binary classification to evaluate the predictive performance of the models and present the results in Table 2 using various metrics. Note that we do not perform any training for this task and use the discretized predictions and ground truth to measure classification performance.

PDSP and MatchMaker outperform other methods in regression performance. PDSP achieves better performances on AUPR (0.81 vs. 0.80) and Precision (0.70 vs. 0.69), MatchMaker is better on F1 (0.72 vs. 0.73) and Recall (0.75 vs. 0.76) metrics.

**Table 2.** Classification performance comparison of the models based on MSE, Pearson and Spearman correlation coefficients.

|  | AUC | pAUC | AUPR | F1 | Precision | Recall | Kappa | BACC |
| --- | --- | --- | --- | --- | --- | --- | --- | --- |
| **PDSP** | 0.79 | 0.64 | 0.81 | 0.72 | 0.70 | 0.75 | 0.40 | 0.70 |
| **MatchMaker** | 0.79 | 0.64 | 0.80 | 0.73 | 0.69 | 0.76 | 0.40 | 0.70 |
| **DeepSynergy** | 0.76 | 0.62 | 0.78 | 0.70 | 0.68 | 0.73 | 0.35 | 0.68 |
| **DeepDDS** | 0.61 | 0.53 | 0.63 | 0.62 | 0.59 | 0.65 | 0.16 | 0.58 |
| **TreeCombo** | 0.73 | 0.60 | 0.75 | 0.68 | 0.67 | 0.79 | 0.32 | 0.66 |
| **Random Forest** | 0.63 | 0.55 | 0.66 | 0.54 | 0.63 | 0.48 | 0.17 | 0.59 |

#### Patient Personalized Classification Performance

While we compare the synergy prediction performance of PDSP with others on cancer cell lines and show above that it is on par with the state-of-the-art methods, its ultimate goal is to perform personalized predictions for cancer patients. Cancer cell line measurements provide large scale training data, but they fail to capture patient variation as cancer is a complex and heterogeneous disease. We have very limited patient-level ground truth data to train a complex machine learning model. For this reason, we employ transfer learning approach in the design of PDSP and use the single drug responses of patients to fine-tune the model as explained in Section 2.4. We compare the classification performances of PDSP (uncalibrated), MatchMaker, DeepSynergy, TreeCombo, DeepDDS and Random Forest models with two patient-calibrated PDSP^st1^ and PDSP^st2^ models. Figure 2 shows the number of *True Positives* (TPs), *True Negatives* (TNs), *False Positives* (FPs) and *False Negatives* (FNs).

**Fig. 2.**
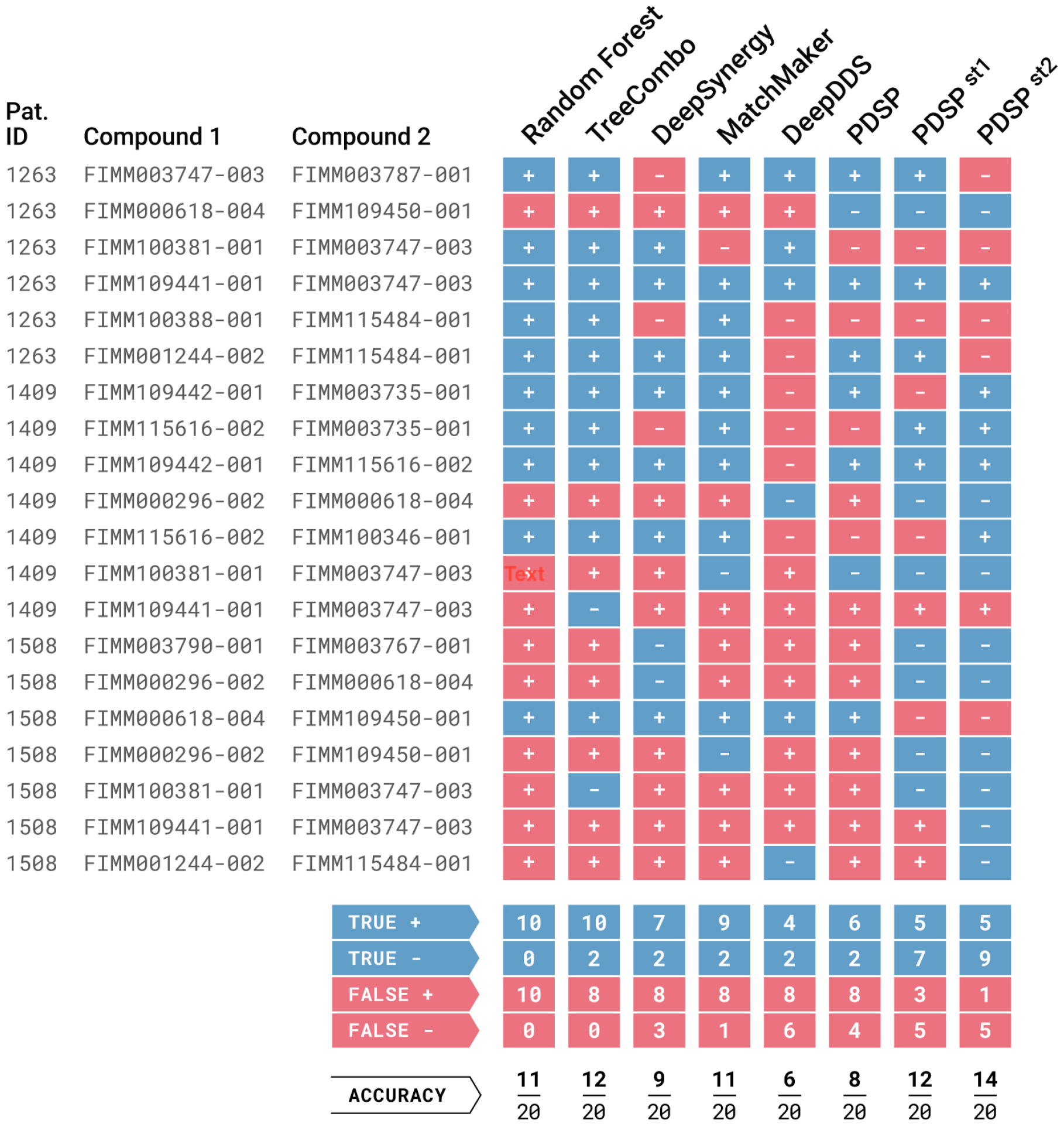
The classification results of patient drug combination synergy data. All models are trained with DrugComb data with cell line gene expressions. PDSP^st1^ and PDSP^st2^ correspond to patient-calibrated models that are fine-tuned based on Strategy 1 and Strategy 2 respectively.

Figure 2 shows that uncalibrated models are not good at detecting antagonistic drug combinations. For example, Random Forest method predicts all drug pair-patient trios as synergistic. Similarly, MatchMaker and TreeCombo predict 17 and 18 (out of 20) drug pairs as synergistic, respectively. They mainly act as a one-class-predictor, which leads to a large number of FPs and low number of TNs. These results lead to low precision values (47% and 56%, respectively) for these models. Compared to uncalibrated models, PDSP^st1^ and PDSP^st2^ achieve better precision (63% and 83%) and separation between synergistic and antagonistic drug combinations. Specifically, the patient-calibrated PDSP^st1^ model decreases the number of FPs to 3 and increases the number of TNs to 7 compared to PDSP. This shows that fine-tuning with patient gene expression improves precision (63% vs. 43%) and accuracy (60% vs. 40%). PDSP^st2^ model performs even better with 9 TNs at the cost of 1 FP. This indicates that it is nearly perfect at detecting antagonistic drug combinations while preserving the performance at synergistic drug combinations. Thus, using fine-tuning and using per-patient calibration (Strategy 2) in particular, yields the best-personalized drug synergy prediction performance.

## 4 Discussion

This study is the first attempt to perform in-silico personalized drug synergy prediction for cancer patients that uses large-scale gene expression data. With limited patient-level data, we opted for a transfer learning based approach to first train a complex model using large scale cell-line based drug-pair perturbation datasets and used patients’ drug response profiles to fine-tune and personalize the model. We show favorable results compared to the state-of-the-art models that do not allow incorporating personal features.

The results indicate that fine-tuning a cell line pre-trained model with a small number of single-drug responses enables PDSP to identify synergistic and antagonistic drug pairs for specific patients better than other models. While the uncalibrated PDSP model’s accuracy is 40% when tested for patients, fine-tuned PDSP^st1^ and PDSP^st2^ models’ accuracies are 60% and 70%, respectively. We observe that uncalibrated models tend to make more synergistic predictions, and they cannot detect antagonistic drug pairs. This problem is alleviated by fine-tuning.

This study uses the drug’s chemical structure and cell line specific gene expression profiles as the feature set. As indicated in [19], these features are selected as they are widely available for most drugs and provide a large training set to train a complex model like ours. If available on a large scale, our model can always be supplemented with other sources of information such as drug-disease or drug-target interactions and network-based methods [7]. Similarly, it is straightforward to adapt the model to use other synergy scores such as Loewe, ZIP and HSA.

We hypothesize that personal single-drug responses can be indirectly used to fine-tune the PDSP model toward better personal drug synergy prediction. We modify the MatchMaker architecture to use this information. We show that, indeed it benefits the drug synergy prediction task in patients. However, as personal drug synergy labels become widely available, we foresee that transfer learning using actual synergy labels will result in even better performance. This is because models will be able to incorporate directly relevant labels, and the model architecture will be less complex with fewer parameters to optimize.

