## Supplementary File for "From Cell-Lines to Cancer Patients: Personalized Drug Synergy Prediction"

### 1 Supplementary Tables

**Table 1.** Top 10 cell line performance for PDSP

| Cell line | # of occurrences in training data | MSE | Pearson | Spearman |
|---|---|---|---|---|
| OVCAR-4 | 2934 | 10.56 | 0.59 | 0.48 |
| SNB-75 | 2822 | 11.56 | 0.65 | 0.49 |
| NCI-H226 | 2991 | 13.37 | 0.60 | 0.48 |
| LOVO | 365 | 13.56 | 0.40 | 0.44 |
| A427 | 365 | 13.70 | 0.68 | 0.66 |
| TK-10 | 2962 | 14.10 | 0.67 | 0.50 |
| UWB1289 | 365 | 14.23 | 0.56 | 0.54 |
| OV90 | 365 | 14.30 | 0.57 | 0.65 |
| EKVX | 2876 | 14.72 | 0.60 | 0.51 |
| OVCAR-5 | 2970 | 14.92 | 0.66 | 0.53 |

**Table 2.** Top 10 cell line performance for MatchMaker

| Cell line | # of occurrences in training data | MSE | Pearson | Spearman |
|---|---|---|---|---|
| OVCAR-4 | 2934 | 10.66 | 0.59 | 0.46 |
| SNB-75 | 2822 | 12.03 | 0.63 | 0.47 |
| LOVO | 365 | 13.25 | 0.38 | 0.43 |
| UWB1289 | 365 | 13.51 | 0.56 | 0.56 |
| NCI-H226 | 2991 | 13.79 | 0.58 | 0.46 |
| TK-10 | 2962 | 14.52 | 0.66 | 0.49 |
| OV90 | 365 | 14.75 | 0.54 | 0.60 |
| A427 | 365 | 15.04 | 0.64 | 0.66 |
| SK-OV-3 | 3301 | 15.68 | 0.62 | 0.52 |
| OVCAR-5 | 2970 | 15.78 | 0.63 | 0.53 |

**Table 3.** Top 10 cell line performance for DeepSynergy

| Cell line | # of occurrences in training data | MSE | Pearson | Spearman |
|---|---|---|---|---|
| OVCAR-4 | 2934 | 12.30 | 0.52 | 0.36 |
| OV90 | 365 | 14.46 | 0.58 | 0.60 |
| LOVO | 365 | 14.67 | 0.28 | 0.31 |
| SNB-75 | 2822 | 14.75 | 0.53 | 0.41 |
| UWB1289 | 365 | 15.47 | 0.52 | 0.53 |
| NCI-H226 | 2991 | 16.70 | 0.48 | 0.37 |
| PC-3 | 2944 | 17.35 | 0.46 | 0.40 |
| A498 | 3059 | 17.50 | 0.54 | 0.44 |
| EKVX | 2876 | 18.33 | 0.51 | 0.41 |
| OVCAR-5 | 2970 | 18.51 | 0.59 | 0.49 |
